# Determining glycosyltransferase functional order via lethality due to accumulated O-antigen intermediates, exemplified with *Shigella flexneri* O-antigen biosynthesis

**DOI:** 10.1101/2023.12.13.571447

**Authors:** Jilong Qin, Yaoqin Hong, Makrina Totsika

## Abstract

The O antigen (OAg) polysaccharide is one of the most diverse surface molecules of Gram-negative bacterial pathogens. The structural classification of OAg based on serological typing and sequence analysis is important in epidemiology and surveillance of outbreaks of bacterial infections. Despite the diverse chemical structures of OAg repeating units (RUs) described, the genetic basis of the assembly of these RUs remains poorly understood and represents a major limitation in our understanding of the gene functions in the polysaccharide biosynthesis. Here we describe a genetic approach to interrogate the functional order of glycosyltransferases (GTs). Using *Shigella flexneri* as a model, we established an initial glycosyltransferase (IT) controlled system to allocate the functional order of the subsequent GT in a two-fold manner. First by reporting the growth defects caused by the sequestration of UndP through disruption of late GTs, and second by comparing the molecular sizes of stalled OAg intermediates when each putative GT is disrupted. Using this approach, we demonstrate that for RfbF and RfbG, the GTs involved in the assembly of *S. flexneri* backbone OAg RU, RfbG is responsible for both the committed step of OAg synthesis, as well as the third transferase for the second L-Rha. We also show that RfbF functions as the last GT to complete the *S. flexneri* OAg RU backbone. We propose that this simple and effective genetic approach can be also extended to define the functional order of enzymatic synthesis of other diverse polysaccharides produced both by Gram-negative and Gram-positive bacteria.

**Importance:** The genetic basis of enzymatic assembly of structurally diverse OAg repeating units in Gram-negative pathogens is poorly understood, representing a major limitation in our understanding of gene functions for the synthesis of bacterial polysaccharides. We present a simple genetic approach to assigning glycosyltransferase (GT) functions and the order in which they act in the assembly of the OAg RU in their native background strain with reasonable confidence. We demonstrated with this approach the functional allocation of GTs for *Shigella flexneri* O antigen assembly. This approach can be generally applied in interrogating GT functions encoded by other bacterial polysaccharides to advance our understanding of diverse gene functions in the biosynthesis of polysaccharides, key knowledge in advancing biosynthetic polysaccharide production.

## Introduction

The O antigen (OAg) is a polysaccharide made of oligosaccharide repeating units (RUs) that forms part of the Gram-negative bacterial surface lipopolysaccharide (LPS). Bacterial OAg is an important virulence factor as its loss renders many Gram-negative bacteria sensitive to host antimicrobials [1, 2] and attenuates colonisation of host niches [3]. Surface OAg structures are also major targets for host niche-residing bacteriophages, and the host immune system due to its high immunogenicity. Moreover, OAg are major glycans directly contributing to host cell attachment through specific glycan: glycan interactions [4], and indirectly contributing to pathogenesis by modulating the exposure of other virulence factors [5, 6], hence constantly subjected to strong selection pressure. These are thought to be evolutionary drivers of OAg diversity, giving rise to over 180 forms of OAg structures characterised so far solely in *E. coli* (including *Shigella* strains) [7], representing one of the most variable bacterial cell surface components. The variability of OAg is due to both the saccharides presented in the OAg RU and the linkages within and between the RU [7] catalysed by a variety of glycosyltransferases (GTs) and polymerases. The variability of O antigenic structures was therefore harnessed to be the basis of a serotyping scheme [8] applied to many Gram-negative bacteria and is of epidemiological importance [9].

Genes for the biogenesis of OAg are clustered in a locus located between the *galF* and *gnd* genes in the majority of *E. coli* genomes [7]. The O locus encodes proteins responsible for a) the biosynthesis of nucleotide diphospho (NDP) sugars as OAg RU precursors, b) GTs for the assembly of OAg RU, and c) RU polymerisation and translocation. Genes for the synthesis of NDP sugars, RU polymerisation and translocation are generally identifiable by sequence alone due to conservation in their primary sequence, secondary structure and protein function based on biochemically well characterised protein families[10]. In contract, the genes for GTs, although often assigned a general GT function by sequence similarity alone to, are difficult to functionally assign to specific sugar linkages with confidence [11, 12]. In some occasions, GT gene function can be predicted with confidence by combining sequence information with information on the OAg chemical structure [2]. However, OAg structures with linkages between multiple saccharides that are identical provides limited guidance in allocating GT functions, representing a major limitation in assigning gene functions for bacterial polysaccharides.

The majority of OAg (93%) in *E. coli* are synthesised by the Wzx/Wzy-dependent pathway (Figure 1A) [7] and have GlcNAc or GalNAc as their initial sugar that is transferred onto the lipid carrier undecaprenol phosphate (UndP/C_55_-P) by the initial GT (IT) WecA [13]. The *wecA* gene is located in the gene cluster for the biosynthesis of enterobacterial common antigen (ECA). The GlcNAc-phosphotransferase reaction by the IT WecA is reported to be reversible (Figure 1A) and the resulting product UndPP-GlcNAc is also the substrate for the second GT (2^nd^ GT) in the ECA biogenesis [14]. In contrast, the glycosyltransferase reaction by the 2^nd^ GT in the OAg biogenesis, which adds the second sugar onto UndPP-GlcNAc is non-reversible and therefore is the committed step in OAg biosynthesis (Figure 1A). Owing to the specificity of GTs for its lipid-linked acceptor, the complete OAg RU is synthesised in a strict sequential manner to be then flipped across the inner membrane (IM) by the flippase Wzx (Figure 1A) [15, 16]. Wzx has specificity towards OAg RU in that only the complete RU is efficiently flipped across the IM (Figure 1A) [17, 18]. The specificity of both GTs and Wzx are the key characteristic in ensuring the correct cell surface presentation of OAg, as the ligase WaaL, which ligates OAg onto the lipid-A core forming LPS, lacks specificity towards its donors [19]. Disruption of OAg sugar precursor biosynthesis, GTs and Wzx will stall the OAg biogenesis [15, 18, 20]. However, there is lack of a feedback mechanism to control OAg biosynthesis when the cytosolic steps beyond the committed step (catalysed by the 2^nd^ GT including the flipping step catalysed by Wzx) are disrupted, as cells with such disruptions accumulate significant amount of UndPP-linked OAg intermediates and suffer severe growth stress creating lethal phenotypes (Figure 1A)[15, 17, 21]. This is due to the sequestration of the universal lipid carrier UndP by the OAg intermediates [17], which needs to be released in the periplasm (PP) by Wzy and WaaL and be recycled efficiently back into cytoplasm (CP) for the biogenesis of other cell envelope components especially for peptidoglycan (PG), an essential synthesis pathway for cell viability [22]. Here we demonstrate that the lack of a feedback-control mechanism could be exploited experimentally as a simple and effective strategy to determine the functional order of GTs in OAg biosynthesis, in particular the 2^nd^ GT in the gene cluster, whereby only the disruption of late GTs as well as Wzx, but not the committed step catalysed by the 2^nd^ GT would accumulate OAg intermediates and generate lethal phenotypes.

**Figure 1.**
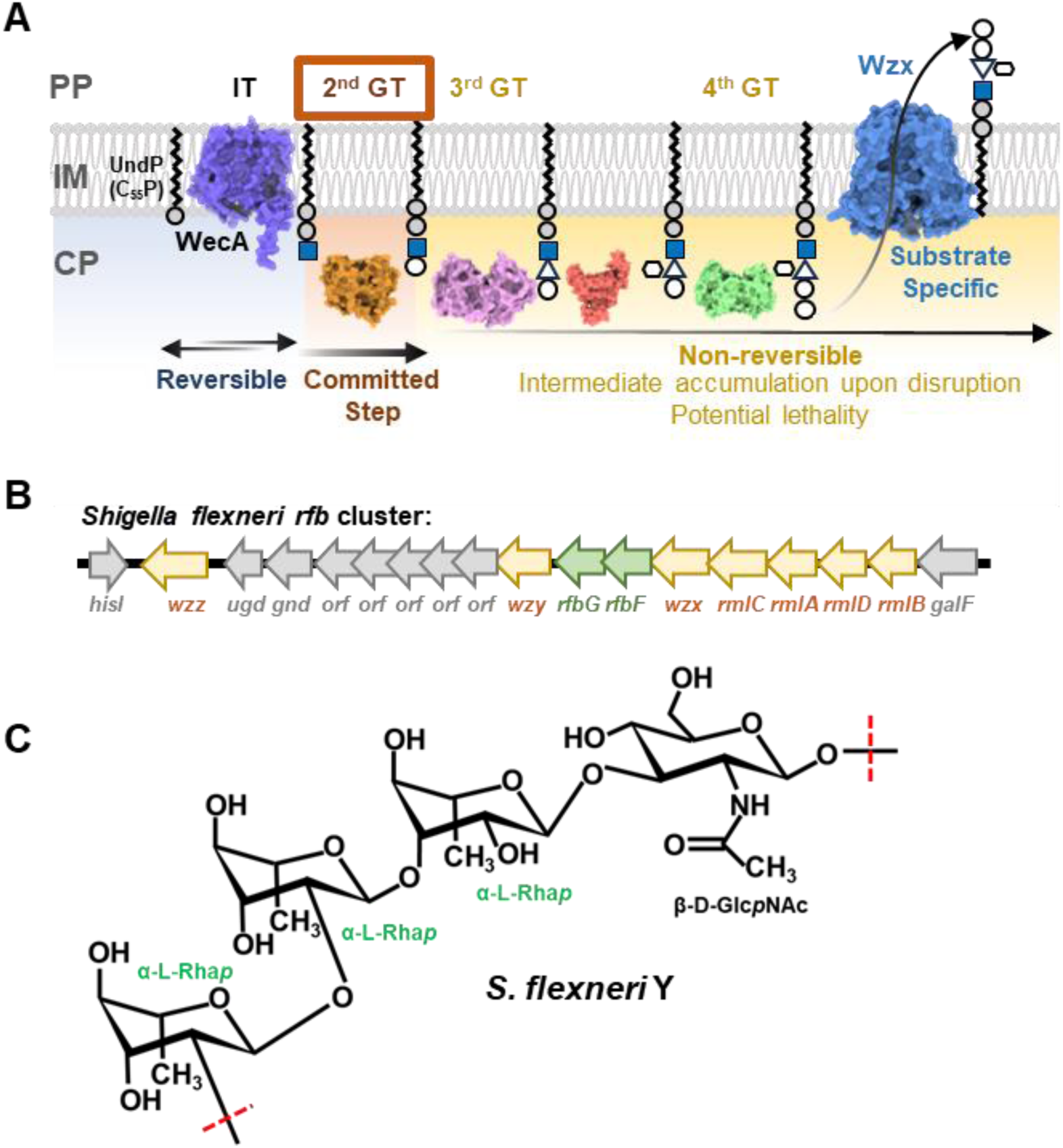
OAg RU biogenesis and *S. flexneri* Y OAg structure. **A)** Stringency of OAg RU synthesis pathways exemplified with K-12 O16 OAg biogenesis. The incomplete OAg RU intermediates on the lipid carrier UndP is not the preferred substrate of Wzx and will remain in the cytosolic face until the complete RU is synthesised. The transfer of saccharide by the second GT is the committed step of OAg biogenesis and disruptions beyond the second GT including Wzx will cause growth defects due sequestration of UndP. PP, periplasm, IM, inner membrane, CP, cytoplasm. **B)** Schematic representation of *S. flexneri rfb* gene cluster. Genes encoding for glycosyltransferases RfbF and RfbG are highlighted in green. **C)** The chemical structure of *S. flexneri* Y serotype OAg.

We use the OAg biosynthesis pathway in *Shigella flexneri* Y serotype as proof of principle of a genetic approach to confidently identify the 2^nd^ GT in the *rfb* locus in a system where the expression of IT is tightly controlled. The *rfb* region of *Shigella flexneri* Y is sufficiently responsible for the synthesis and assembly of *Shigella flexneri* Y serotype OAg [23] (Figure 1B), which has the chemically defined RU structure →2)-α-L-Rha*p*III-(1→2)-α-L-Rha*p*II-(1→3)-α-L-Rha*p*I-(1→3)-β-D-Glc*p*NAc-(1→ [24] (Figure 1C) and is shared by all *S. flexneri* serotype strains as the backbone structure. However, only two genes (*rfbF* and *rfbG,* (referred as *wbgF* and *wbgG*, respectively, according to the bacterial polysaccharide gene nomenclature (BPGN) scheme [25, 26]) are predicted to be rhamnose glycosyltransferases in the *rfb* region (Figure 1B). It is therefore difficult to allocate RfbF and RfbG functions in *S. flexneri* Y OAg sugar linkages. We demonstrate that disruption of RfbF as well as Wzx, while not RfbG, generated lethal phenotypes, and confirm the role of RfbG as the 2^nd^ GT in *S. flexneri* OAg synthesis. By comparing the molecular sizes of the small dead-end intermediates that were capped onto LPS molecules in strains with lethality, we were also able to determine that RfbG is responsible for the transfer of the first two rhamnoses, while RfbF is responsible for the last rhamnose. This is the first study to assign OAg GT functions in *S. flexneri*, demonstrated as an example for the allocation of functional order for polysaccharides GTs via a genetic approach.

## Results

### RfbG is the second GT in the assembly of S. flexneri Y OAg

As a first step we deleted *wecA* encoding for the OAg IT in *S. flexneri* strain PE860, and abolished the production of Smooth LPS (S-LPS) in this strain (Figure 2A). This was done to avoid potential deleterious effects of subsequent mutations introduced in the OAg late GTs, which may lead to genetic instability and secondary suppressor mutations that would confound the interpretation of results in our study. We then engineered a WecA expression construct under *pBAD* promoter control, the expression of which could complement the *ΔwecA* mutant to produce S-LPS similar to that of WT PE860 upon arabinose induction (Figure 2A). We firstly confirmed that the induction of WecA in the *ΔwecA* mutant background didn’t not have any observable effect on cell viability (Figure 2B) or growth kinetics (Figure 2C). We then successfully deleted the *wzx* flippase gene in the *ΔwecA* background with no impact on cell viability. In contrast, initiation of OAg RU synthesis by induction of WecA in *ΔwecAΔwzx* severely affected cell viability (Figure 2B) and caused cells lysis when growing in liquid media (Figure 2C). This is consistent with the results reported for a *Salmonella enterica ΔgalEΔwzx* mutant with OAg production under exogenous galactose control, showing that Wzx is essential when the OAg RU is assembled [21], confirming that the sequestration of UndP by OAg dead-end intermediates affects cell viability and growth, creating observable lethal phenotypes in systems with OAg production under tight control. We then deleted the two GTs *rfbF* and *rfbG*, respectively, in the *ΔwecA* background, and both were confirmed to have no observable effect in cell viability (Figure 2B). However, when OAg production was induced by *in trans* WecA expression, the *ΔwecAΔrfbF* mutant but not the *ΔwecAΔrfbG* showed severe defects in cell viability and increased cell lysis (Figure 2B-C), similar to that observed with *ΔwecAΔwzx* mutant expressing WecA. Disruption of GTs acting directly after WecA in OAg assembly process will prevent UndP entering disrupted OAg synthesis pathways and abolish the sequestering effect, as the product of WecA UndPP-GlacNAc will be redirected into making ECA or be reverted into UndP by WecA as discussed previously. Therefore, our results suggest that i) RfbG is the 2^nd^ GT catalysing the committed step for *S. flexneri* Y OAg RU assembly; and ii) RfbF acts after RfbG, and when disrupted, leads to accumulated intermediates that are not efficiently recognised and flipped by Wzx, thereby sequestering UndP in the OAg synthesis pathway and creating lethal phenotypes.

**Figure 2.**
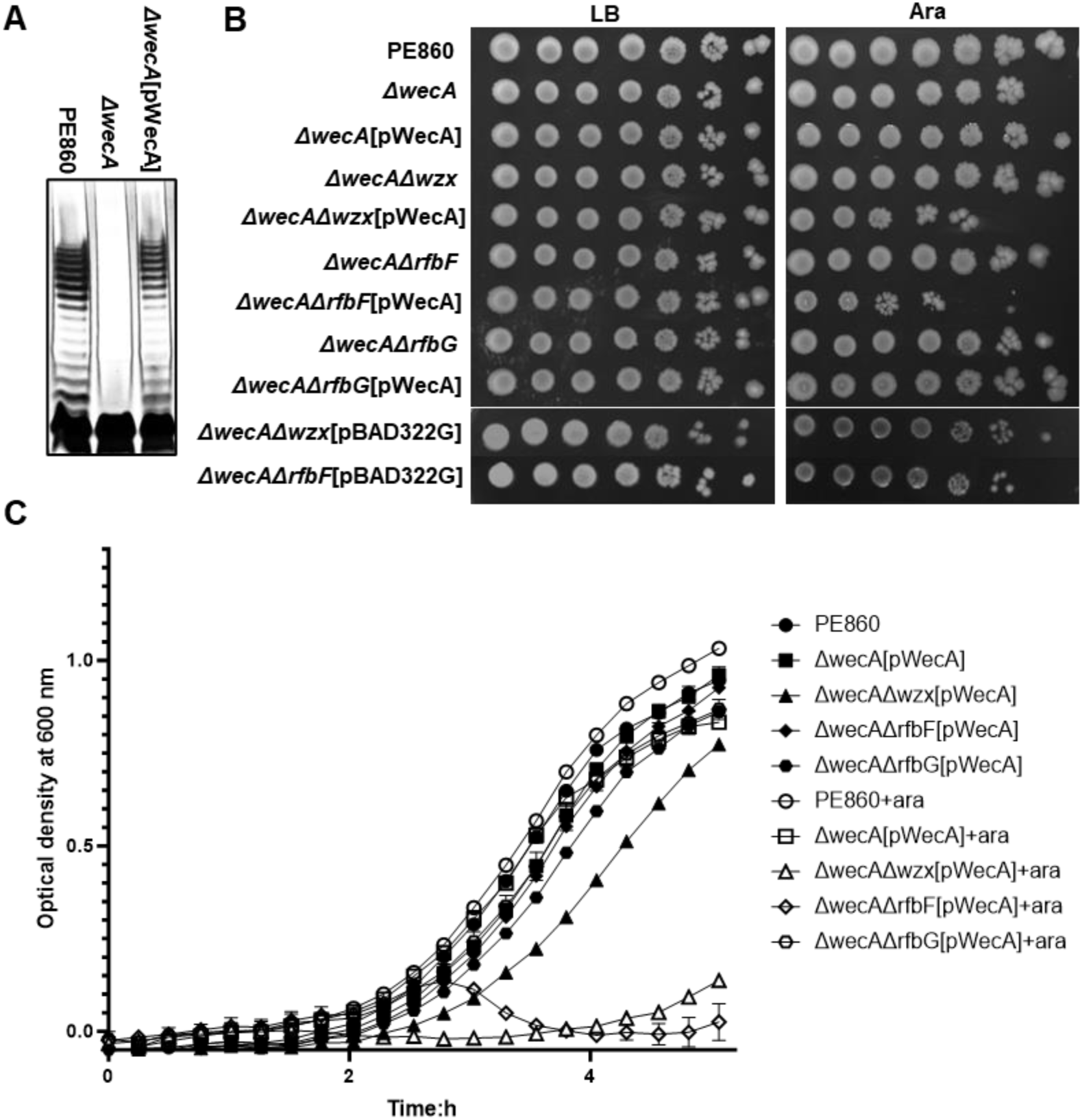
Lethality caused by OAg intermediate accumulation in *S. flexneri* Y serotype strains with disruptions in late GTs and translocase Wzx. **A)** LPS silver staining of samples prepared from indicated *S. flexneri* strains. **B)** Bacterial cultures of indicated strains grown in LB were adjusted to OD_600_ of 1 and 10-fold serial dilutions (10^0^ to 10^−6^) were spotted (4 μl) onto LB agar supplemented without or with 10 mM L-arabinose (ara). **C)** Growth curves of indicated *S. flenxeri* strains in LB without or with 10 mM arabinose (ara) for pWecA induction.

### RfbG cross-complements E. coli K-12 producing O16 OAg

1. *E. coli* K-12 strain MG1655 is devoid of its O antigen due to a characterised mutation in *wbbL* [27] which encodes the 2^nd^ GT responsible for the formation of L-Rha*p*(1→3) linkage in the O16 OAg backbone structure (→2)-β-D-Galf-(1→6)-α-D-Glcp-(1→3)-α-L-Rhap-(1→3)-α-D-GlcpNAc-(1→) (Figure 3A) [28]. The L-Rha*p*(1→3) linkage between O16 and *S. flexneri* Y (Figure 1C) is identical, hence to validate the GT function of RfbG, we expressed both RfbG and RfbF in MG1655, respectively. Indeed, expression of RfbG but not RfbF restored the production of S-LPS in MG1655 (Figure 3B), albeit at reduced level in comparison to the *wbbL* repaired MG1655-S strain [20]. Western immunoblotting with monospecific anti-O16 antiserum confirmed that the restored S-LPS is compose of O16 OAg (Figure 3C). Therefore, our results confirmed the role of RfbG in acting as the 2^nd^ GT catalysing L-Rha*p*(1→3) linkage in the *S. flexneri* Y OAg.

### RfbG is a dual L-Rha transferase and RfbF is the last GT for S. flexneri Y OAg synthesis

Since RfbG could functionally complement the *wbbL* mutation in *E. coli* K-12 restoring O16 OAg production, to further assign the function of RfbG and RfbF to the glycosidic linkages between the remaining two L-Rha in *S. flexneri* Y OAg, we then cross-complemented the PE860*ΔwecAΔRfbG* mutant with plasmids expressing WecA and WbbL under the tight control of *pBAD* and *pTet* promoters, respectively. Interestingly, while induction of either WecA or WbbL alone in PE860*ΔwecAΔRfbG* showed no growth impact in liquid media, induction of both simultaneously caused cell lysis (Figure 4A). This suggests that expression of both WecA and WbbL in PE860*ΔwecAΔRfbG* caused accumulation of OAg intermediates that could not be recognised and processed by the downstream RfbF, thereby sequestering UndP and triggering lethality.

**Figure 3.**
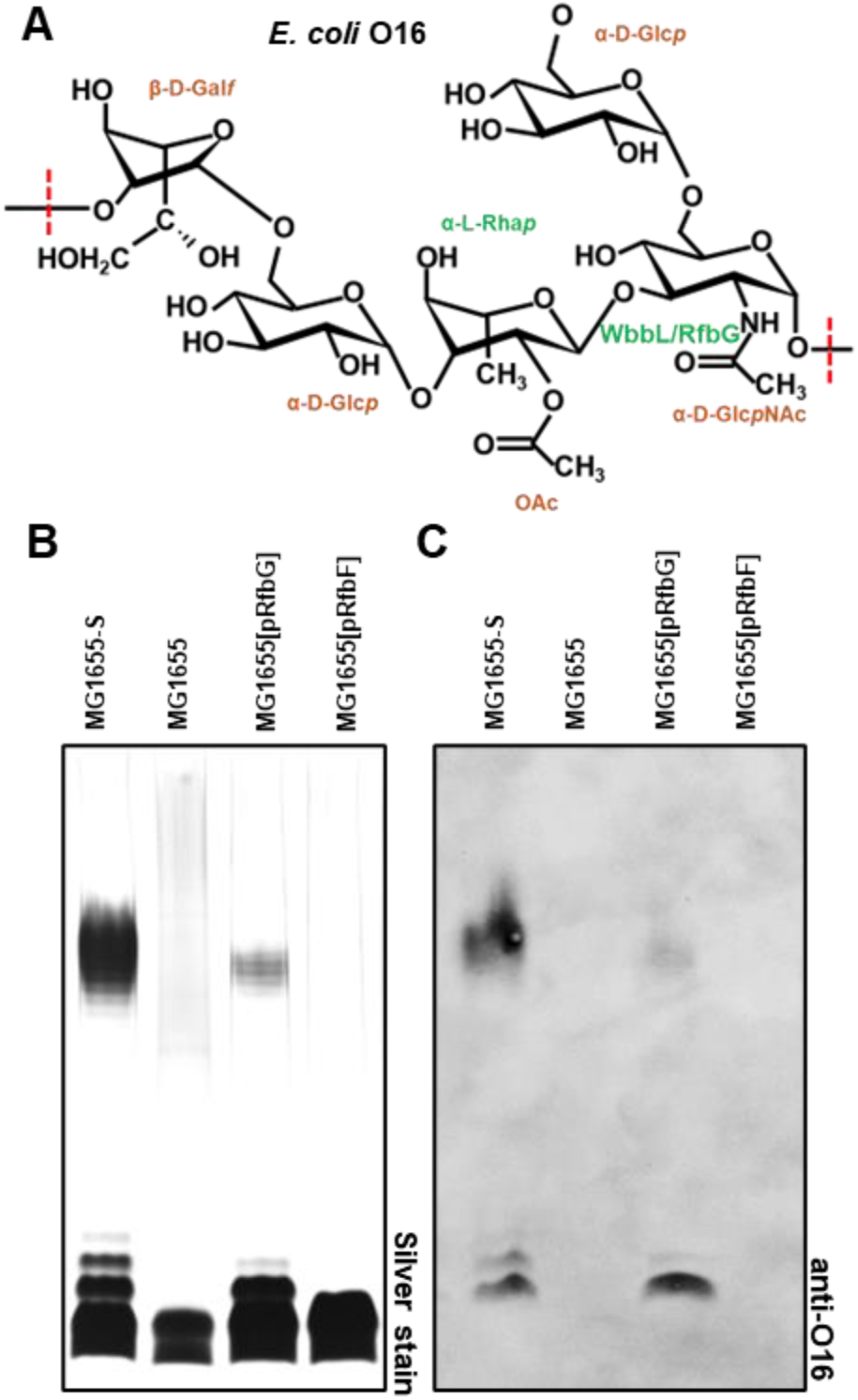
Cross-complementation of *E. coli* K-12 O16 O-antigen production by *S. flexneri* glycosyltransferases. **A)** Same glycosidic linkage catalysed by WbbL and RfbG shown with *E. coli* K-12 O16 O antigen structure. **B)** LPS silver staining of *E. coli* K-12 MG1655 complemented with *S. flexneri* glycosyltransferases. **C)** Western immunoblotting of samples of *S. flexneri* strains shown in **B**) with anti-O16 antibodies.

**Figure 4.**
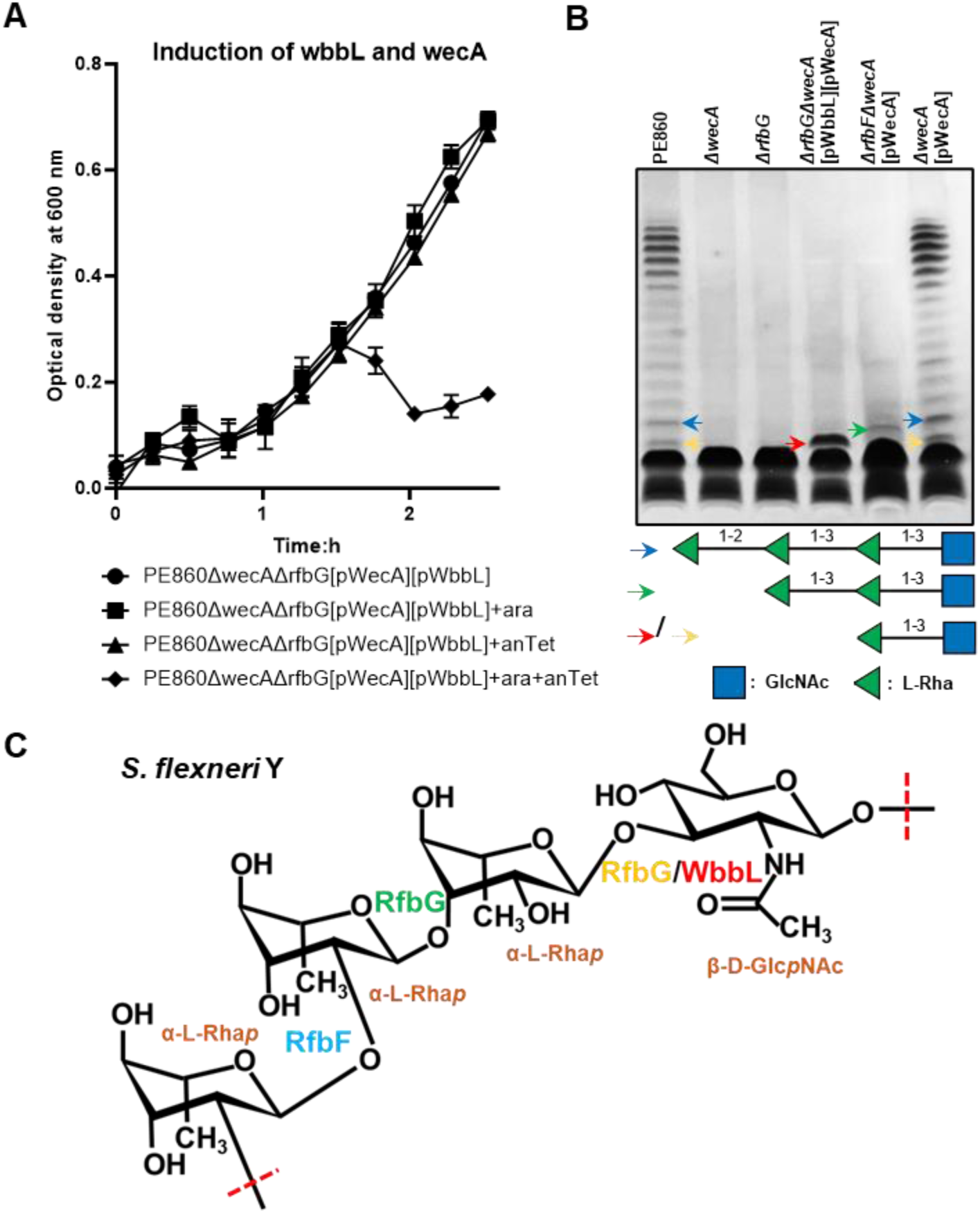
Ligation of incomplete *S. flexneri* O-antigen intermediates onto LPS in cells with lysis. **A)** Growth curves of indicated *S. flexneri* strains with the expression of WecA non-induced or induced with 10 mM arabinose (ara), and/or WbbL non-induced or induced with 50 ng/ml anhydrotetracycline (anTet). **B)** LPS silver staining of samples of indicated *S. flexneri* strains with early lysis. The predicted O-antigen saccharides capped onto LPS are marked by arrows and corresponding structures shown (GlcNAc-Rha in yellow and red, GlcNAc-Rha-Rha in green, and GlcNAc-Rha-Rha-Rha in blue). **C)** Proposed glycosidic linkages catalysed by RfbG and RfbF shown in the *S. flexneri* Y O antigen structure.

The accumulation of lipid-linked OAg intermediates in the cytoplasmic side of IM is due to the specificity of Wzx, leading to deficient translocation across the physical barrier to reach WaaL in the periplasmic side of IM and be ligated onto Lipid-A core to form semi-rough LPS. However, we hypothesised that strains showing the cell lysis phenotype could also have a disrupted IM and small amounts of intermediates gaining access to WaaL to form semi-rough LPS. Indeed, by analysing with silver staining the LPS of bacterial samples harvested at the beginning of cell lysis, we were able to compare the molecular sizes of these OAg intermediates by SDS-PAGE (Figure 4B). In the absence of *wecA* or *rfbG*, only the band of lipid A-core can be detected (Figure 4B), suggesting that the UndPP-GlcNAc was not sufficiently flipped by the Wzx of PE860. The expression of both WecA and WbbL in PE860*ΔwecAΔRfbG* showed a semi-rough LPS (Figure 4B, red arrow) band in relatively high amount. The molecular size of this semi-rough molecule is larger than the R-LPS band shown in both PE860*ΔwecA* and PE860*ΔrfbG* but is smaller than the semi-rough LPS band of the PE860*ΔwecAΔRfbF* expressing WecA (Figure 4B, green arrow). The two semi-rough LPS bands detected in PE860*ΔwecAΔRfbG* expressing WecA and WbbL and PE860*ΔwecAΔRfbF* expressing WecA were both smaller than the LPS with 1 RU of the WT PE860 or PE860*ΔwecA* expressing WecA (Figure 4B, blue arrow), hence are the lipid A-core capped with different OAg intermediates. These data infer that the semi-rough LPS band detected in PE860*ΔwecAΔRfbF* expressing WecA is lipid A-core-GlcNAc-Rha-Rha, and that the PE860*ΔwecAΔRfbG* expressing both WecA and WbbL produced lipid A-core-GlcNAc-Rha. Interestingly, in the WT PE860 (Figure 2A & Figure 4B) and PE860*ΔwecA* expressing WecA (Figure 4B), a semi-rough band having the same molecular size of that detected in PE860*ΔwecAΔRfbG* expressing both WecA and WbbL was consistently detected, suggesting that Wzx of *S. flexneri* Y could also inefficiently recognise and flip UndPP-GlcNAc-Rha across IM. Taken together, we propose that RfbG is both the 2^nd^ and 3^rd^ GT transferring the first two L-rhamnoses onto UndPP-GlcNAc, and RfbF is the last GT transferring the third L-rhamnose to synthesise the complete *S. flexneri* Y OAg RU (Figure 4C).

## Discussion

The OAg in the LPS is highly immunogenic and is a target of humoral responses in the host. The highly diverse structures of OAg between and within species provides the basis for serological typing of Gram-negative bacterial pathogens and is important for epidemiology and surveillance of emerging pathogens. Despite advancements in OAg typing of pathogens via serology or more recently whole genome sequence data analysis in conjunction with over 180 solved OAg structures [7], determination of the genetic basis of RU assembly for the synthesis of different O antigen structures remains a challenging task and represents a major hurdle in our understanding of gene functions for polysaccharide biosynthesis. This is because the numerous combinations of different donor sugars (at least 20 different sugars may compose the OAg), acceptor sugars/oligosaccharides, sugar anomerism, yet there is lack of biochemical evidence to characterise all GT families found so far as both donor sugar in their NDP activated form and the lipid linked incomplete OAg RU are not readily available. Therefore, it is rarely possible to assign GTs’ function and predict the order in which they may act to assemble the RU structure with confidence through sequence analysis alone. Our study provides a simple method to effectively determine the GT functional order in the OAg RU assembly, which enabled us to successfully allocate the functions of RfbG and RfbF in the assembly of *S. flexneri* backbone OAg. Firstly, by comparing growth after mutating each of the putative GT genes in a system where the IT expression is tightly controlled, the committed step (2^nd^ GT) of OAg RU assembly can be identified by having no growth impact when mutated. Secondly, in conjunction with the available chemical structure of the OAg RU, comparisons of the mass of the accumulated intermediates after mutating GTs individually allows to allocate the function and the order of each GT on OAg RU assembly with confidence.

Our genetic approach for determining the committed GT function relies on the tight control of IT, a key strategy to avoid genetic instability when constructing GT deletion mutants. This is because the accumulation of dead-end intermediates via direct deletion of late GTs generates lethal conditions, potentially allowing a selection of suppressors preventing UndP entering the disrupted polysaccharide assembly pathways [29, 30]. These phenomena may alter or complicate the interpretation of results. Therefore, we propose that when studying GT functions with late GT mutants, a parent strain with controlled IT or GT at the committed step expression should be used.

Disruption of the OAg production is a common strategy in constructing live attenuated vaccines. Our approach in determining the lethal phenotypes when deleting late GTs in the polysaccharide’s assembly pathways hence could provide important guidance in the construction of attenuated vaccine strains devoid of polysaccharide, in that the direct late GT mutations should be avoided to ensure the genetic stability of the live vaccine strain. It is interesting that a live attenuated *Shigella*-enterotoxigenic *E. coli* (ETEC) vaccine was constructed by direct deletion of *rfbF* [31]. However, our results here suggests that this mutant vaccine strain may contain secondary mutations preventing UndP in making OAg during strain construction or the *rfbF* mutation in the vaccine strain is polar to the downstream *rfbG*. In addition, OAg represents an important vaccine antigen itself when coupled with protein carrier through bioconjugation, known as glycoconjugates [32], which was demonstrated to be effective in animals against a range of bacterial pathogens with some proceeded to phase 2 clinical trials [33]. The successful production of desired bacterial polysaccharides heterologously in the engineered vaccine-producing host strain requires knowledge in understanding the gene functions in the glycan assembly, as it can be difficult to predict the OAg produced in strains carrying heterologous *rfb* genes. Indeed, introduction of *S. flexneri rfb* locus with *Tn* insertions in *rfbF* into *E. coli* K-12 was reported previously to produce an altered OAg type different to S *flexneri* Y serotype [11]. With the GT gene functions assigned for *S. flexneri rfb* region in this work, we predicted and confirmed that the altered OAg produced in this strain is O16 due to the functional complementation of *wbbL::IS5* by *rfbG*. Therefore, our approach paves the way in expanding the knowledge required for the genetic construction of glycoconjugate vaccine strains.

Determination of OAg GT functions in *E. coli* using a genetic approach has been reported previously [34]. In the previous work, a tester strain was developed with all OAg, ECA and colanic acid synthesis genes deleted to eliminate any potential heterologous sugar structure modifications, and with the IT expression under tight control. The *rfb* regions from the target strain with putative GTs were cloned in a plasmid and introduced into the tester strain, allowing the analysis of the mass of the intermediates to infer GT functions. However, this approach requires substantial work in the construction of the tester strain and the assembly of *rfb* locus (∼10 kb) of the target strain into a plasmid. In addition, the approach solely relies on the analysis of the mass of the intermediates when GTs were mutated. However, the copy number of the *wzx* genes introduced into the tester strain via plasmid could influence the OAg intermediate-linked LPS produced. This is because increased Wzx expression could have phenotypically lessened specificity [17], potentially accommodating earlier steps of OAg precursors that otherwise not stalled in the synthesis to be flipped across the IM, complicating the interpretation of results. In contrast, our approach of GT function determination was done in the parent strain background, with no changes in gene copies and expression levels for *wzx*.

A feedback mechanism in controlling the assembly of OAg RU in the cytosolic face of IM beyond the committed step is lacking, building dead-end intermediates when late GT were mutated triggering lethal phenotype [21]. Harnessing this knowledge, we innovatively demonstrated a simple genetic approach by just comparing the lethal phenotypes of mutants with each putative GT mutated to determine the 2^nd^ GT function. The lack of the feedback mechanism in the assembly of RUs of polysaccharides seems to exist more widely and beyond OAg synthesis pathways, as shown for the ECA biosynthesis pathway [35], and capsular polysaccharide synthesis pathways in other Gram-negative bacteria [36, 37] and Gram-positive bacteria [30, 38]; the latter also representing diverse major polysaccharide antigens important for epidemiology and clinical serology. Therefore, our genetic approach for distinguishing the committed steps of putative GT between the late GTs, in principle can be extended also for other diverse polysaccharide type as well as to other bacteria strains including Gram-positive bacteria.

## Materials and Methods

### Bacterial strains and plasmids

The bacterial strains and plasmids used in this work are listed in Table 1. Single colonies of bacterial strains grown overnight on Lysogeny Broth (LB)-Lennox [39] agar (1.5% w/v) plates were picked and grown overnight in LB at 37 °C for subsequent experiments. Where appropriate, media were supplemented with ampicillin (Amp, 100 µg/ml), kanamycin (Kan, 50 µg/ml), chloramphenicol (Chl, 25 µg/ml), anhydrotetracycline (AhTet 10 ng/ml), or arabinose (Ara, 10 mM).

**Table 1.**
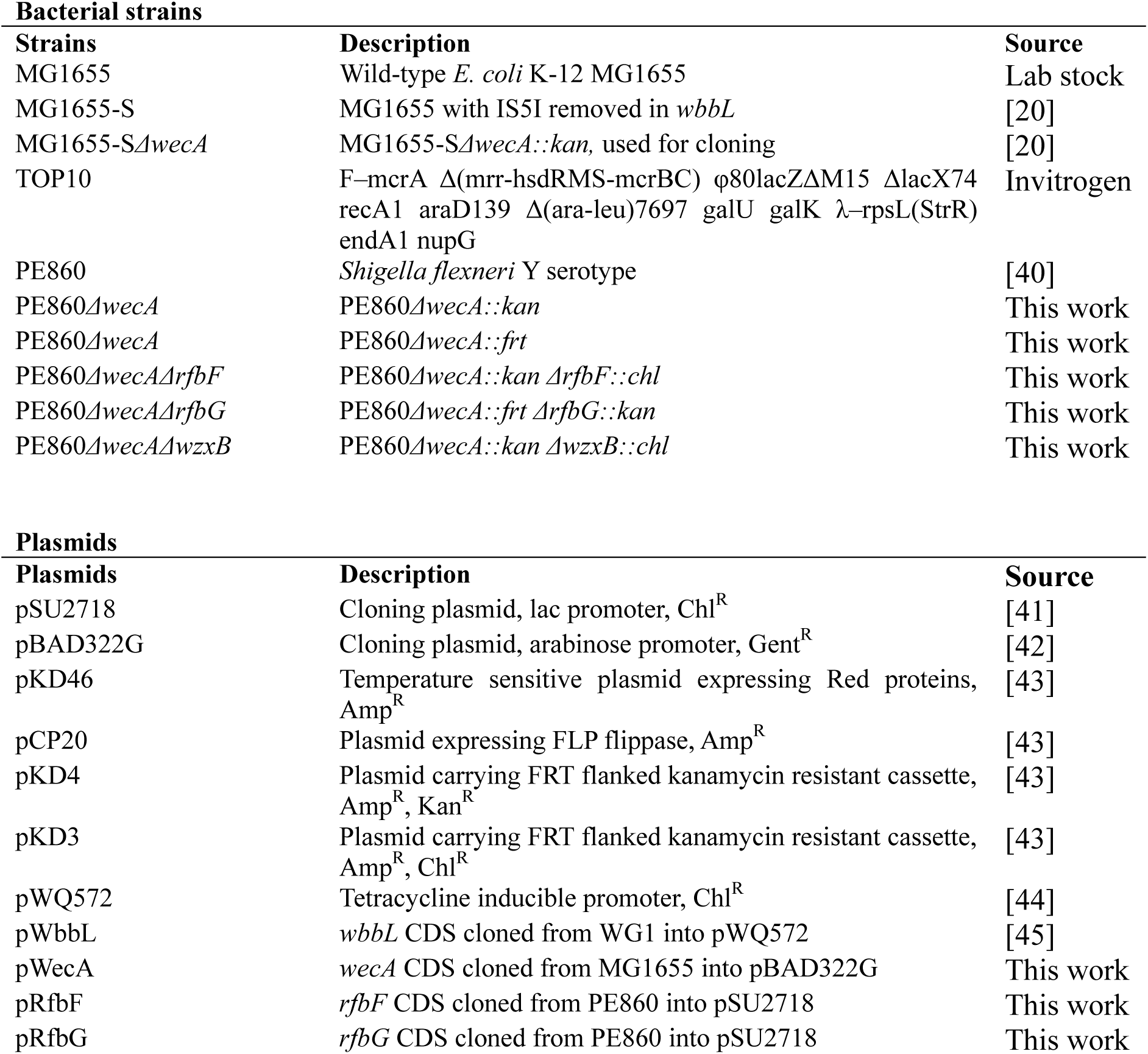
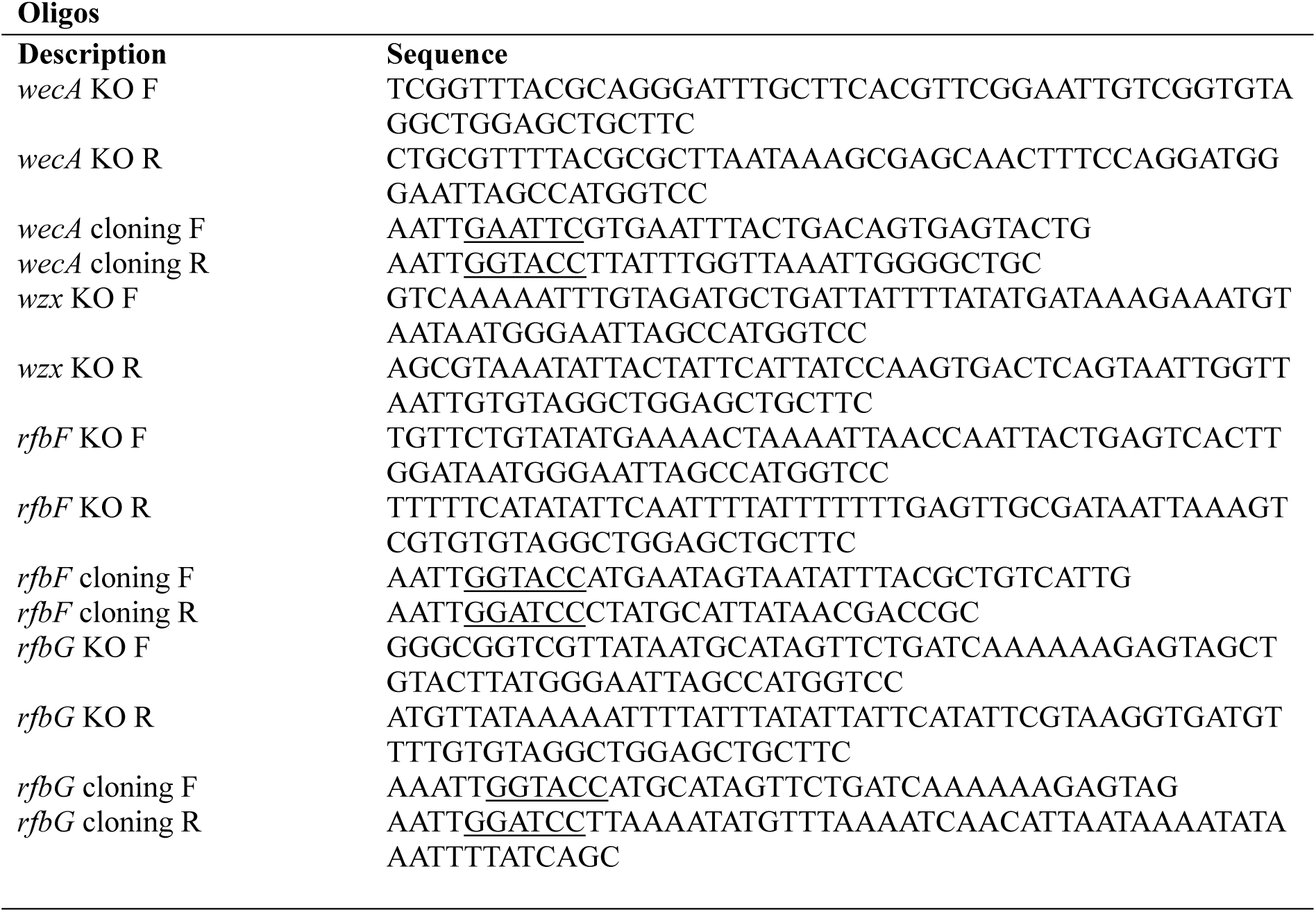
Strains, plasmids, and oligonucleotides.

### Bacterial mutagenesis via allelic exchange

Mutagenesis was performed as described previously [43] with laboratory adapted optimisations [46]. Briefly *Shigella flexneri* strains harbouring plasmid pKD46 grown overnight in 10 ml LB at 30 °C was sub-cultured 1 in 100 into 10 ml LB in a 50 ml tube. Expression of the lambda phage-derived Red proteins were then induced with 50 mM L-arabinose at OD_600_ of 0.3 for 1 h. Bacterial cells were then harvested via centrifugation, washed twice with 10 ml ice-cold water, and resuspended in 100 μl of 10% (v/v) ice-cold glycerol for subsequent electroporation. The *cat* or *neo* gene was PCR amplified from pKD3 or PKD4, respectively, with primers containing 50 bp up- and down-stream sequences homologous to the target gene (Table 1). The PCR amplicon was purified (1.5 μg) and introduced into electrocompetent cells via electroporation and cells were immediately recovered in 3 ml LB in a 50 ml Falcon tube that was incubated for 2 h at 37 °C before plating out (100 μl) on LB agar plates supplemented with Chl or Kan. Plates were incubated at 37 °C for 16 h to acquire mutants. Successful mutants were PCR screened and confirmed.

### Plasmid construction

For generation of expression constructs, the coding sequences of WecA, RfbF and RfbG were PCR amplified from boiled whole-cell water preparation of PE860 and cloned into pBAD322G (for WecA) and pSU2718 (for RfbF and RfbG) using restriction enzyme cloning, resulting in pWecA, pRfbF and pRfbG, respectively. RfbF and RfbG clones were recovered from MG1655-S*ΔwecA* to avoid potential O antigen intermediate build up in cloning strains.

### Bacterial survival spotting assay

Bacterial survival spotting assay were performed as described previously [20]. Briefly, overnight bacterial cultures were adjusted to optical density at 600 nm (OD_600_) of 1 and 10-fold serial diluted to 10^-7^ with fresh LB media. A 4 μl of each dilution preparations were spotted onto LB agar plates supplemented with or without 10 mM arabinose.

### Bacterial growth kinetic assay

Bacterial growth kinetic were recorded as described previously [20]. Briefly, overnight bacterial cultures were diluted 1 in 200 μl of fresh LB media, supplemented with or without 10 mM arabinose and/or 10 ng/ml anhydrotetracycline in a 96-well plate. Plates were incubated at 37°C with aeration in a CLARIOstar plate reader (BMG, Australia) programmed to measure the OD_600_ every 6 minutes over 18 h.

### LPS silver staining

LPS silver staining were performed as described previously[20]. Briefly, bacterial cells (10^9^) grown at mid-exponential phase were collected via centrifugation (20,000 g 1 min) and lysed in 50 μl of SDS sample buffer and heated at 100°C for 10 min. Samples were then cooled and treated with 50 μg/ml proteinase K (PK, NEB) for 18 h at 60°C. PK-treated samples were then heated at 100°C for 10 min, and 2-5 μl sample were loaded onto 10-20% SDS-tricine gels (Invitrogen, #EC66252BOX) and LPS was silver stained as described previously [47]. For bacterial strains producing O-antigen intermediates substituted LPS, cells were grown to OD_600_ of 0.8 in LB with 0.2% (w/v) glucose, followed by washing with fresh LB media 2 times, then induced with 10 mM arabinose and/or 50 ng/ml anhydrotetracycline for 20 min to allow early cell lysis to occur. LPS samples were then prepared and analysed as above.

### Western immunoblotting

For Western immunoblotting of O16 LPS, polysaccharide samples separated by SDS-tricine gel electrophoresis were transferred onto nitrocellulose membrane and detected with rabbit polyclonal anti-O16 antibodies (SSI Diagnostica, #SSI85012).

## Acknowledgement

The authors thank A/Prof. Renato Morona (The University of Adelaide) in providing *Shigella flexneri* Y strain PE860.

## Financial statements

This work is funded in part by an Australian Research Council project grant (DP210101317), the Max Planck Queensland Centre on the Materials Science of Extracellular Matrices to MT, and an Early Career Research Ideas Grant from the Faculty of Health, Queensland University of Technology (Australia) to JQ. The Ian Potter Foundation sponsored the CLARIOStar high-performance microplate reader (BMG, Australia). The funders had no role in study design, data collection and analysis, decision to publish, or preparation of the manuscript.

## Author contributions

JQ concepsulised the project; JQ and YH contributed to experimental design; JQ conducted experiments, and contributed to data collection, analysis and interpretation; JQ and MT supervised the study and obtained the funding. JQ wrote the manuscript and all authors edited the manuscript.

## Competing interests

The authors declare no competing interests.

